# Interactions between social groups of colobus monkeys (*Colobus vellerosus*) explain similarities in their gut microbiomes

**DOI:** 10.1101/717934

**Authors:** Eva Wikberg, Diana Christie, Pascale Sicotte, Nelson Ting

## Abstract

The gut microbiome is structured by social groups in a variety of host taxa. Whether this pattern is driven by relatedness, similar diets, or shared social environments is under debate because few studies have had access to the data necessary to disentangle these factors in wild populations. We investigated whether diet, relatedness, or the 1-meter proximity network best explains differences in the gut microbiome among 45 female colobus monkeys in 8 social groups residing at Boabeng-Fiema, Ghana. We combined demographic and behavioural data collected May-August 2007 and October 2008-April 2009 with 16S rRNA sequencing of faecal samples collected during the latter part of each observation period. Social group identity explained a large percentage of the variation in gut microbiome beta-diversity. When comparing the predictive power of dietary dissimilarity, relatedness, and connectedness in the 1-meter proximity network, the models with social connectedness received the strongest support, even in our analyses that excluded within-group dyads. This novel finding indicates that microbes may be transmitted during intergroup encounters, which could occur either indirectly via shared environments or directly via social contact. Lastly, some of the gut microbial taxa that appear to be transmitted via 1-meter proximity are associated with digestion of plant material, but further research is needed to investigate whether this type of gut microbe transmission yields health benefits, which could provide an incentive for the formation and maintenance of social bonds within and between social groups.

## Introduction

The gut microbiome consists of thousands of species that affects its host’s nutritional status, immune function, and behavior (McFall-Ngai et al., 2013). It is associated with parasite resistance and stress response of hosts in the wild (Koch & Schmid-Hempel, 2011; Vlčková et al., 2018) and with obesity in captive settings (Turnbaugh et al., 2006). Because of these potential health consequences, it is important to investigate the acquisition and maintenance of the gut microbiome (Amato, 2016; Archie & Tung, 2015).

The gut microbiome of individuals or social groups become more distinct with geographic distance (Barelli et al., 2015; Grieneisen et al., 2019; Hansen et al., 2019; Hird, Carstens, Cardiff, Dittmann, & Brumfield, 2014; Lankau, Hong, & Mackie, 2012; Phillips et al., 2012), and the microbiome is structured by social group or family co-residency in a variety of host taxa, such as humans (Lax et al., 2014; Song et al., 2013; Yatsunenko et al., 2012), non-human primates (Amato et al., 2017; Degnan et al., 2012; Goodfellow et al., 2019; Orkin, Webb, & Melin, 2019; Springer et al., 2017; Tung et al., 2015), carnivores (Leclaire, Nielsen, & Drea, 2014; Theis et al., 2013), birds (White et al., 2010), and insects (Anderson et al., 2012; Koch & Schmid-Hempel, 2011). For example, the gut microbiome of immigrant male baboons *(Papio cynocephalus)* converge over time with that of their new group members (Grieneisen, Livermore, Alberts, Tung, & Archie, 2017), and the gut microbiomes of colobus monkeys (*Colobus vellerosus*) diverged over the course of nine months after a social group fissioned into two daughter groups (Goodfellow et al., 2019).

The divergence in gut microbiomes with home range separation could potentially be due to dietary differences, lower degrees of relatedness, or lack of shared social environments (Archie & Tung, 2015; Björk, Dasari, Grieneisen, & Archie, 2019). Diet is suggested to be one of the most important factors affecting the gut microbiome (Voreades, Kozil, & Weir, 2014). Gut microbial composition fluctuates within hosts with seasonal or experimental dietary changes (Davenport et al., 2014; David et al., 2013; Hicks et al., 2018; Leamy et al., 2014; Mallott, Amato, Garber, & Malhi, 2018; Maurice et al., 2015; Michl et al., 2019; Orkin, Campos, et al., 2019), and dietary similarities may explain whether social groups have distinct gut microbiomes (Orkin, Webb, et al., 2019). However, this pattern could also reflect the genetic similarity of hosts in societies where at least some closely related individuals remain together in their natal group. When this is the case, closely related group members are expected to have more similar gut microbiomes than non-group members with lower degree of relatedness, because the host’s genetic makeup affects microbe colonization (Opstal & Bordenstein, 2015; Spor, Koren, & Ley, 2011) and a number of genomic regions are associated with gut microbial composition in rodents (Bonder et al., 2016; Leamy et al., 2014). This may explain why closely related individuals have more similar gut microbiomes than unrelated individuals in some studies of humans and captive rodents (Faith et al., 2013; Kovacs et al., 2011; Ley et al., 2005). In contrast, genetic differentiation between baboon populations was a poor predictor of their gut microbiome (Grieneisen et al., 2019), and relatedness did not have a significant effect on the gut microbiome in some studies of humans (Rothschild et al., 2018), non-human primates (Moeller et al., 2016) and carnivores (Leclaire et al., 2014). Moeller and colleagues (2016) suggest that this may be due to an overriding effect of transmission among unrelated social partners. Indirect social contact via shared environments, such as touching common surfaces, may facilitate microbiome transmission within households (i.e., indirect social transmission) (Lax et al., 2014). Direct social contact such as grooming or sitting in body contact further increases microbiome transmission (i.e., direct social transmission) between close social partners within social groups of monkeys (*Alouatta pigra*: Amato et al., 2017; *Papio cynocephalus*: Tung et al., 2015; Grieneisen et al., 2017) and lemurs *(Eulemur rubriventer:* Raulo et al., 2017). Gut microbiomes are also more similar among socially connected than disconnected siblings and married couples (Dill-McFarland et al., 2019). In contrast, social connectedness between non-group members did not predict gut microbiome similarity in sifakas (*Propithecus verreauxi*) (Perofsky et al., 2017). Even if intergroup encoutners promote the transmission of microbes, there may not be an association between intergroup interactiowns and gut microbiome similarity if groups rarely interact at close distances. Taken together, these studies indicate that gut microbes are transmitted via social interactions within social groups, while it is unclear whether this is the case for social interactions between social groups.

To investigate whether the pattern of increasing between-individual differences in the gut microbiome (i.e., beta-diversity) with home range separation is best explained by lower dietary overlap, relatedness, or social connectedness, we focus on the black-and-white colobus monkeys (*Colobus vellerosus*) at Boabeng-Fiema, Ghana. This is one of several rare species of arboreal leaf-eating monkeys distributed across the forested regions of the African tropics, and it is closely related to guerezas (*Colobus guereza*) and western black-and-white colobus (*Colobus polykomos*) (Ting, 2008). At Boabeng-Fiema, all colobus social groups utilize a highly folivorous diet, but the most important food species differ between social groups (Saj & Sicotte, 2007; Teichroeb & Sicotte, 2009). More seeds and fruits are available during the dry season, during which they eat up to 43% of these food items (Teichroeb & Sicotte, 2017). To break down hard-to-digest items in their primarily folivorous diet (Saj & Sicotte, 2007; Teichroeb & Sicotte, 2009), they rely on behavioural traits, physiological traits, and their gut microbiome (Amato et al., 2016; Lambert, 1998). Possibly due to constraints imposed by their highly folivorous diet, colobus monkeys spend a low percentage of their time engaging in direct social activities such as grooming (Teichroeb, Saj, Paterson, & Sicotte, 2003). Female colobus spend on average 3% of their time within 1 meter and 0.1% of their time grooming each female group member (Wikberg, Ting, & Sicotte, 2014b). However, females still form preferred friendships, which are only occassionally based on kinship and never based on their relatively weakly expressed dominance hierarchies (Wikberg, Teichroeb, Bădescu, & Sicotte, 2013; Wikberg, Ting, & Sicotte, 2014a; Wikberg et al., 2014b; Wikberg, Ting, & Sicotte, 2015). Instead, females prefer to affiliate with females with similar immigration status (Wikberg et al., 2014b, 2014a) in this population where all males and half of the females disperse (Sicotte et al., 2017; Teichroeb, Wikberg, & Sicotte, 2009, 2011; Wikberg, Sicotte, Campos, & Ting, 2012). This flexible female dispersal pattern results in social groups with different female kin composition and some close maternal female kin residing in different social groups (Wikberg et al., 2012). Neighbouring social groups encounter each other in the large zones of home range overlap on an almost daily basis. During these encounters, social groups sometimes chase each other away from food trees, while at other times, they engage in affiliative or sexual between-group interactions (Sicotte & MacIntosh, 2004; Teichroeb & Sicotte, 2017).

The frequent between-group interactions coupled with variation in diet and relatedness within and between social groups makes this a good study population to investigate whether the pattern of increasing gut microbial beta-diversity with home range separation is best explained by lower degrees of dietary similarity, relatedness, or social connectedness. We take a cross-sectional approach using observational and genetic data from eight social groups to first test whether the gut microbiome was structured by social groups. We predicted gut microbiome beta-diversity to be structured by social groups and to increase with home range separation. We then evaluated which factors explained gut microbiome beta-diversity between females across different social groups. We expected gut microbiome beta-diversity to decrease with dietary similarity and relatedness and increase with distance in the 1-meter proximity network. Finally, the significant predictor from the analyses above (social connectedness) was used in a subsequent population-level analysis of Operational Taxonomic Unit (OTU) abundance to determine which microbial taxa may be socially transmitted. Our definition of social transmission includes both direct social transmission via physical contact and indirect social transmission via shared substrates (e.g., Perofsky et al., 2017), and we will not attempt here to tease these two social transmission routes. Males and females of all age-classes were used to create social networks, but the gut microbiome data are only available for adult females. Therefore, our analyses of beta-diversity focus on adult females.

## Methods

### Behavioral data collection

Demographic data have been collected since 2000 from the black-and-white colobus monkeys (*Colobus vellerosus*) at Boabeng-Fiema, Ghana. In this study, we also use behavioral and ecological data as well as DNA samples from eight social groups (Fig. A1) collected during two study periods: the rainy season May-August 2007 and the pre-dry and dry seasons October 2008 - April 2009 (Table A2). During this time period, the study groups contained 3-9 adult (i.e., parous) females (Table A2), 1-4 adult males, and 8-24 immatures. Our research adheres to ASAB/ABS Guidelines for the Use of Animals in Research, the laws of Ghana, and data collection was approved by the Boabeng-Fiema Monkey Sanctuary’s management committee, Ghana Wildlife Division, and the University of Calgary’s Animal Care Committee (BI 2006-28, BI 2009-25).

We recorded the social group’s location every hour using a map with trails, roads, villages, and large trees (>40 cm DBH) in order to determine home ranges (Fig. A1). During 10-minute focal samples (Altmann, 1974) of adult females, we continuously recorded all social behaviors (including the identity of the interactant and the duration of the behavior) and plant species and part (i.e., mature leaf, young leaf, flower, fruit, seed, or other) for each ingested food item. Females fed on a total of 210 food item-plant species combinations, and to assess dietary differences, we calculated Sørensen dissimilarity indices using ingested plant parts and plant species during focal samples. We choose this diversity index because it only takes the presence or absence of an ingested food item into account, which we have a robust estimate of using the focal data. The Sørensen dissimilarity indices in our data set had a high median value of 0.83 and it was lower within than between social groups (Fig. A2).

We observed 61 and 285 between-group encounters (i.e., two social groups located within 50 meters of each other) during the first and second data collection period respectively. Of these encounters, 53% lacked female aggression and 35% lacked male aggression. Because close proximity between individuals of different social groups are rare and unlikely to be recorded during focal sampling, we recorded approaches to 1 meter *ad libitum* (Altmann, 1974). Some of these approaches only led to brief close proximity while others led to prolonged contact like copulations, grooming, and play. We created an undirected proximity network based on the presence and absence of approaches to 1 meter between all individuals (N = 177 adult females, adult males, and immatures) present in the eight study groups. We used the software UCINET (Borgatti, Everett, & Freeman, 2002) to compute inverse shortest path length (i.e., Geodesic distance) in the 1-meter proximity network (hereafter referred to as social connectedness): 1/[the number of steps (i.e., recorded interaction ties) in the shortest path from one individual to another]. Social group members were in 1-meter proximity with each other (i.e., an inverse path length of 1) or separated by two to three partners (i.e., an inverse path length of 0.5 and 0.33) (Fig. A2). The inverse path length for males and females belonging to different social groups ranged from 0 to 1 (Fig. 1; Fig. A2). The seemingly unconnected individuals in the 2007 data set were most likely unconnected because we only had access to data collected from a 3-month period. These individuals were connected and separated with up to eight steps in the 2008-2009 network, which was based on six months of data.

**Figure 1.**
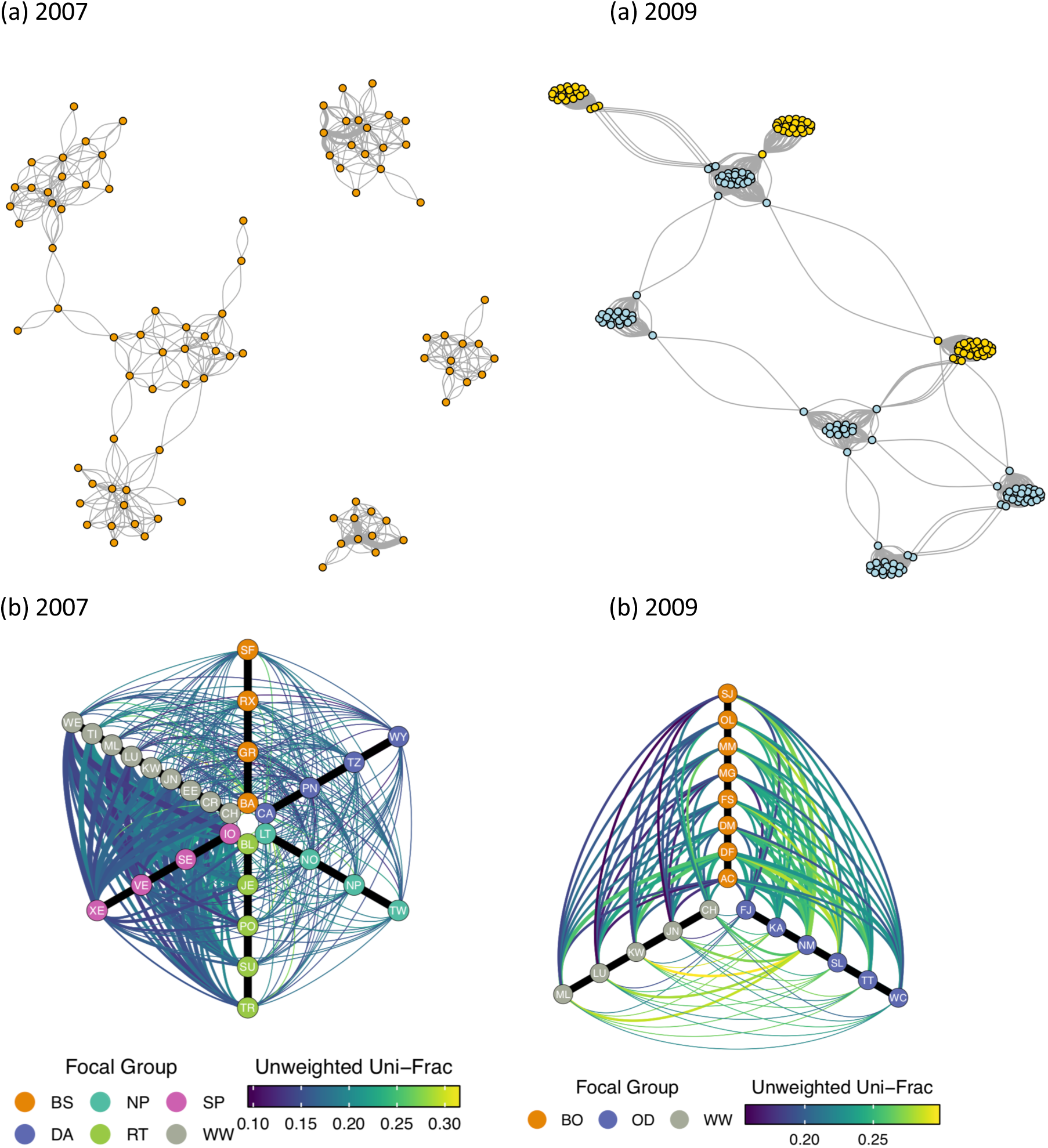
Social networks for: **a)** the entire population where each group member is depicted as a node in yellow (group used for behavioural analyses) or blue (group not used for behavioural analyses) and between-group dyads observed in 1-meter proximity are connected with lines; **b)** females included in the behavioural analyses with lines connecting between-group dyads (i.e., nodes of different color) where color represents gut microbiome beta-diversity (i.e., unweighted Unifrac distances) ranging from similar (dark) to dissimilar (light) and thickness indicates social connectedness ranging from strongly connected (thick) to more disconnected (thin). The black lines connect group members and are not weighted based on beta-diversity or social connectedness.

### Genetic data collection

We collected faecal samples June-August 2007 and January-April 2009. Immediately after a female defecated, we collected approximatey 1g of feces and dissolved it in 6ml RNAlater. The samples were stored in a fridge at the field site until the end of the field season when they were transported to the Ting lab and stored in a −20-degree C freezer. Note that we lack information on soil type, which was driving between-site differences in the gut microbiomes in a large-scale study of terrestrial baboons (Grieneisen et al., 2019). However, our samples were collected from arboreal primates within a small study area, and sampling site does not have a significant effect on beta-diversity in our study population (Goodfellow et al., 2019).

We extracted DNA from the samples and genotyped the extracts at 17 short tandem repeat loci (STR) as previously described (Wikberg et al., 2012). To make sure that the samples used in the relatedness and gut microbiome analyses were collected from the correct individual, we compared the STR genotypes obtained from these samples with a second sample collected from the same individual at a different time. We calculated dyadic estimated relatedness values (*R*) in MLRelate (Kalinowski, Wagner, & Taper, 2006) because this method provided the most accurate relatedness estimates in our study population (Wikberg et al., 2012). We used *R*-values calculated from STR loci rather than theoretical relatedness (*r*) calculated from pedigrees, because *R*-values predict kinship relatively accurately in our study population (Wikberg et al., 2014a) and they are more accurate than *r* in studies such as ours with limited access to pedigrees (Forstmeier, Schielzeth, Mueller, Ellegren, & Kempenaers, 2012; Robinson, Simmons, & Kennington, 2013). The median female relatedness was low both within and between social groups, but there were at least some closely related females residing in the same social groups (Fig. A2).

For generating the gut microbial data, we conducted fresh DNA extracts from 61 previously genotyped samples from 45 females (Table A2) using the QIAamp DNA Stool Mini Kit with a modified protocol. More details regarding the extraction protocol are presented in the Appendix and in Goodfellow et al. (2019). The V4 hypervariable region of the bacterial 16S ribosomal RNA gene was amplified and libraries were prepared using the 515F and 806R primers containing 5’ Illumina adapter tails and dual indexing barcodes, and libraries were sequenced as part of a 150bp paired-end sequencing run on the Illumina NextSeq platform following Goodfellow et al. (2019). We obtained a mean read depth of 127,628 per sample (range: 86,924-166,438). Then, we used a custom pipeline (https://github.com/kstagaman/Process_16S) for quality filtering and assembly (see Appendix). We performed *de novo* OTU picking in UCLUST (Edgar, 2010), and sequences with 97% overlap were defined as belonging to the same bacterial Operational Taxonomic Unit (OTU). After this processing, we had a total of 2,597 OTUs and an average of 89,483 reads per sample (range: 59,817-120,119). To further guard against sequencing errors, we filtered out OTU’s with a frequency lower than 0.00005 as recommended (Bokulich et al., 2012). After filtering, the 2007 data set contained 450 OTUs and the 2009 data set contained 396 OTUs. The mean read depth was 88,346 (range: 59,005 – 118,633). We did not rarefy the data set to an even read depth, because it is recommended against (McMurdie & Holmes, 2014). First, rarefying leads to increased false positives and decreased true positives, especially in data sets with read depths comparable to ours (Pereira et al., 2018). Second, unrarefied counts are particularly accurate when using our measure of beta-diversity—weighted UniFrac distances (McMurdie & Holmes, 2014).

We initially calculated four different measures of gut microbiome beta-diversity (Sørensen dissimilarity index, Bray-Curtis dissimilarity index, unweighted UniFrac distances, and weighted UniFrac distances) in the R package vegan (Oksanen et al., 2017). Because the two presence/absence indices were strongly correlated with each other (Sorenson dissimilarity indices and unweighted UniFrac distances: Mantel r = 0.93, p = 0.001) as were the two abundance indices (Bray-Curtis dissimilarity indices and weighted UniFrac distances: Mantel r = 0.77, p = 0.001), in our analyses, we only used the one presence/absence index (unweighted UniFrac distances) and the one abundance index (weighted UniFrac distances) that take phylogenetic relationships of OTUs into account.

### Data analyses

We combined the 2007 and 2009 data sets and included study year (aka season) and individual ID as predictor variables whenever possible (i.e., permutational multivariate analysis of variance and linear mixed models) while we had to create squared interaction matrices for each study year separately when using matrix correlations (i.e., Mantel tests and Moran’s test for autospatial correlations). We only used the full data set (N = 61 samples from 45 females) for the initial analysis regarding the effect of social group identity. All subsequent analyses examined the effects of behavioural variables on beta-diversity in a subset (N = 49 samples from 42 unique females) from which we removed: 1) duplicate samples from the same year and same female; 2) one adult female with incomplete dietary information, and 3) social groups from which the majority of females remained unsampled to make sure we had a representative sample of social connectedness from each social group.

The initial analysis investigated the effects of season, social group, individual identity, and read depth on beta-diversity of all dyads in the full data set (N = 61 samples from 45 females in 2007 and 2009) using permutational multivariate analysis of variance (PERMANOVA) with 10,000 permutations using the adonis function in the R package vegan (Oksanen et al., 2017). The terms were added sequentially in the order listed above.

We used non-parametric Mantel correlations implemented in the R package vegan (Oksanen et al., 2017) to investigate whether the two measures of gut microbiome beta-diversity were correlated with home range separation (0 = same social group and home range; 1 = different social groups but adjacent home ranges, 2 = different social groups and non-adjacent home ranges) using beta-diversity indices from 30 samples from unique females in 6 social groups in 2007 and 19 samples from unique females in 3 social groups in 2009. We used beta-diversity indices of all dyads, but analysed the two years separately.

To investigate which combination of dyadic traits predicted gut microbiome beta-diversity between females, we created generalized linear mixed models (GLMMs) with the outcome variable gut microbiome beta-diversity using the beta family function in the package glmmTMB (Magnusson et al., 2019) in R (R Core Team, 2018). Again, we used 30 samples from unique females in 6 social groups in 2007 and 19 samples from unique females in 3 social groups in 2009. We created a null model that did not contain any fixed effects, alternative models with one fixed effect that represented one of the hypotheses outlined in the introduction (dietary dissimilarities*, R*-values, or social connectedness), and a full model with all three predictor variables. We included data collection year as a fixed effect in all alternative models because the two sampling years occurred in different seasons and several other studies show strong seasonal shifts in gut microbiome composition (Amato et al., 2015; Hicks et al., 2018; Orkin, Campos, et al., 2019; Smits et al., 2017; Springer et al., 2017). All numerical predictor variables were centered and scaled (Schielzeth, 2010). We included social group and focal identities as random effects in all GLMMs, including the null models. We did not have any issues with collinearity based on low Variance Inflation Factors for the full models (all VIF < 1.43). We evaluated the support for each model using Akaike Information Criterion (AIC) (Akaike, 1974), and this approach allowed us to determine which hypotheses (diet, relatedness, or social connectedness) was best supported by our data (Burnham & Anderson, 2002). Because several models received similar support, we took model selection uncertainly into account by averaging coefficients across models (Burnham & Anderson, 2002) using the R package MUMIN (Barton, 2013). In the first set of analyses, we included dyads that resided in the same social group and dyads that resided in different social groups. To make sure that the effect of social connectedness was not driven by the close social bonds within social groups, we repeated the analyses with between-group dyads only.

To infer which of the gut microbial taxa may be transmitted via close proximity, which was a better predictor of beta-diversity than diet and relatedness (see Results), we investigated whether the abundance of each OTU was correlated with Geodesic distance in the 1-meter approach network using Moran’s test for autospatial correlations implemented in the package ape (Paradis, Claude, & Strimmer, 2004). We included within-group and between-group dyads in this analysis (N = 342 dyads). We counted the number of OTUs in each phylum (or family) that were socially structured based on the autospatial correlation results. We conducted hypergeometric tests to investigate whether this number was higher than expected by chance based on the total number of OTUs in the phylum (or family) using the phyper function implemented in R. In all analyses of taxonomic differences, we used the 10% false discovery rate to correct p-values for multiple testing (sensu Tung et al., 2015). The gut microbial taxa we expected to be shaped by sociality are listed in the Table A1 (Amato et al., 2017; Goodfellow et al., 2019; Tung et al., 2015).

## Results

### Factors predicting gut microbiome beta-diversity

We investigated the relative effects of season, social group, individual ID, and read depth in the full data set (PERMANOVA: N = 61 samples collected 2007-2009). Of the observed variation in the taxonomic composition of the gut microbiome (i.e., beta-diversity), individual identity explained the largest percentage (54-55% depending on which beta-diversity index was used as outcome variable), social group identity explained a more moderate percentage (19-28%), while year explained much smaller percentage (8-12%) (Table 1). Read depth did not have a significant effect on beta-diversity (Table 1).

**Table 1.** Results from the PERMANOVA with factors added sequentially in the ordered listed in the table.

| Beta-diversity index | Factor | Df | Sums of squares | Mean squares | F | R<sup>2</sup> | P |
| --- | --- | --- | --- | --- | --- | --- | --- |
| Unweighted UniFrac | Season | 1 | 0.103 | 0.103 | 12.325 | 0.084 | <0.001 |
|  | Group | 7 | 0.347 | 0.050 | 5.943 | 0.282 | <0.001 |
|  | ID | 39 | 0.663 | 0.017 | 2.035 | 0.538 | <0.001 |
|  | Read depth | 1 | 0.010 | 0.010 | 1.185 | 0.008 | 0.249 |
| Weighted UniFrac | Season | 1 | 0.136 | 0.136 | 12.277 | 0.118 | <0.001 |
|  | Group | 7 | 0.219 | 0.031 | 2.812 | 0.189 | <0.001 |
|  | ID | 39 | 0.639 | 0.016 | 1.477 | 0.553 | <0.001 |
|  | Read depth | 1 | 0.018 | 0.018 | 1.637 | 0.016 | 0.104 |

Gut microbiome beta-diversity and home range separation were correlated in the 2007 data set (N = 870 dyads in 6 social groups, Mantel tests: unweighted UniFrac distance: r = 0.22, P = 0.002; weighted UniFrac distance: r = 0.10, P = 0.049) and in the 2009 data set (N = 342 dyads in 3 social groups, Mantel tests: unweighted UniFrac distance: r = 0.36, P = 0.005; weighted UniFrac distance: r = 0.20, P = 0.024), meaning that females residing farther from each other had less similar gut microbiomes. This pattern can potentially be explained by group members having more similar diets, higher relatedness, or stronger social connectedness than non-group members (Fig. A2).

We created several competing generalized linear mixed models to investigate which of the three hypotheses best explained increasing beta-diversity with home range separation: dietary dissimilarity, relatedness, or social connectedness, controlling for data collection year. In our data set with both within-group and between-group dyads (N = 1,212 dyads in 2007-2009), the full models and the models with social connectedness received the greatest support (Table 2). Social connectedness predicted gut microbiome beta-diversity, and females located further apart in the social network had less similar gut microbiomes (Figs. 1-2). Year also predicted gut microbiome beta-diversity (Fig. 2), and females had more similar gut microbiomes during the rainy season of 2007 than the dry season of 2009. In contrast, diet and relatedness did not have significant effects on gut microbiome beta-diversity (Fig. 2).

**Figure 2.**
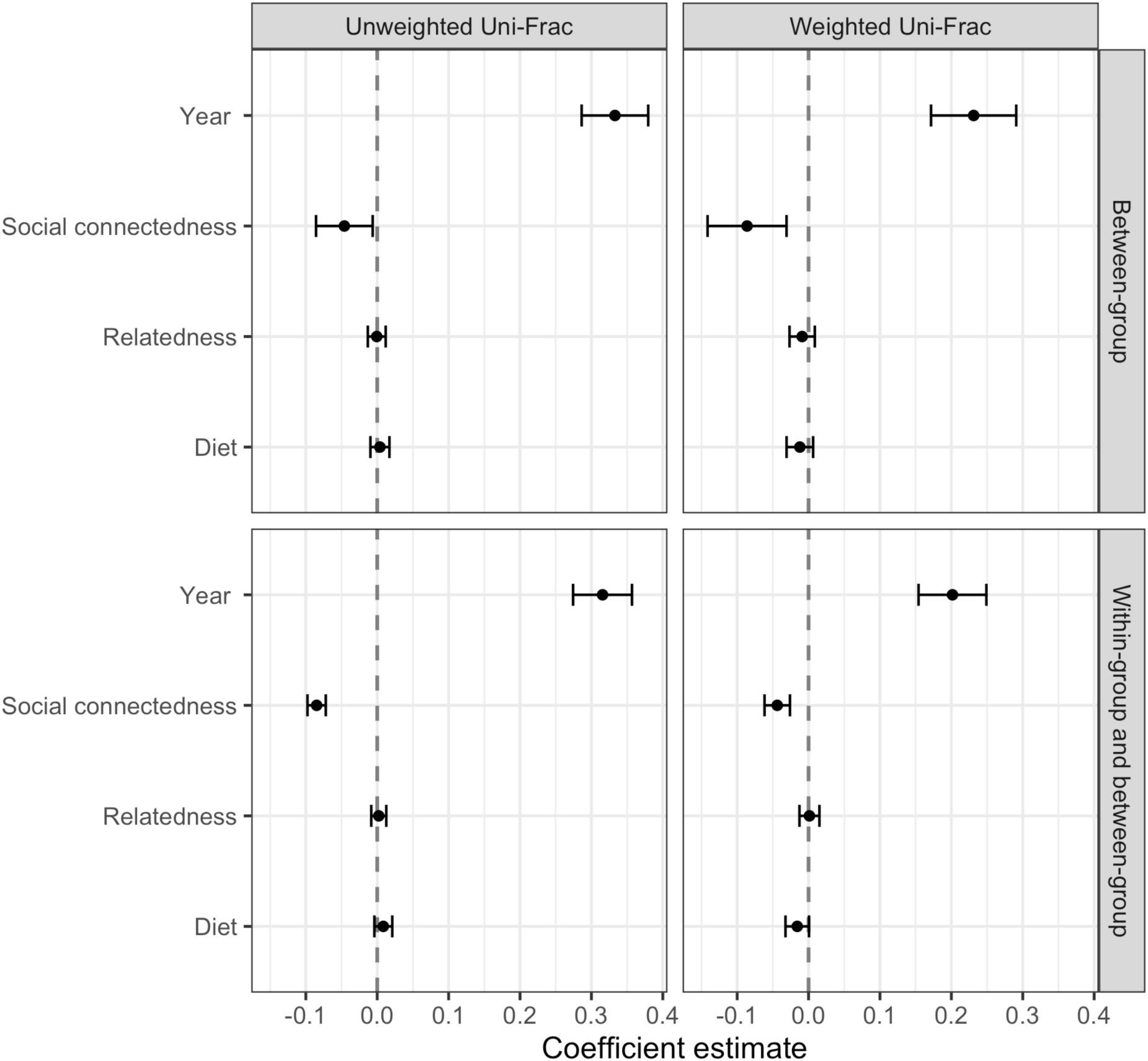
Coefficient estimates and their 95% confidence intervals for the best fitting model for unweighted versus weighted UniFrac distances in the data set including within-group and between-group dyads and in the data set with between-group dyads only. Social connectedness, dietary dissimilarity, and relatedness ranges from 0 to 1. Of the two study periods, year 2007 was used as the baseline level against which we depict the effect of year 2009.

**Table 2.** The competing GLMMs’ fixed effects, Akaike Information Criterion, delta (i.e., difference in AIC between the current model and the best-fit model), and Akaike weights (i.e., relative likelihood of the model), and marginal and conditional R^2^ for the best fitting model (i.e., without versus with random effects) when including **a)** within-group and between-group dyads and **b)** only between-group dyads.

| Outcome variable | Fixed effect | AIC | Delta | Weight |
| --- | --- | --- | --- | --- |
| **Unweighted UniFrac** | Season + Proximity | -5247.30 | 0.00 | 0.73 |
|  | Season + Diet + Relatedness + Proximity | -5245.28 | 2.02 | 0.27 |
|  | Season + Diet | -5103.63 | 143.67 | 0.00 |
|  | Season + Relatedness | -5060.36 | 186.94 | 0.00 |
|  | - | -4940.73 | 306.58 | 0.00 |
| **Weighted UniFrac** | Season + Proximity | -4684.20 | 0.00 | 0.56 |
|  | Season + Relatedness | -4683.69 | 0.51 | 0.44 |
|  | Season + Diet | -4658.94 | 25.25 | 0.00 |
|  | Season + Diet + Relatedness + Proximity | -4657.85 | 26.35 | 0.00 |
|  | - | -4614.36 | 69.84 | 0.00 |
| **Unweighted UniFrac** | Season + Proximity | -4307.62 | 0.00 | 0.76 |
|  | Season + Diet | -4303.92 | 3.71 | 0.12 |
|  | Season + Relatedness | -4302.67 | 4.95 | 0.06 |
|  | Season + Diet + Relatedness + Proximity | -4302.44 | 5.19 | 0.06 |
|  | - | -4119.55 | 188.07 | 0.00 |
| **Weighted UniFrac** | Season + Proximity | -3770.21 | 0.00 | 0.64 |
|  | Season + Diet | -3768.91 | 1.30 | 0.33 |
|  | Season + Relatedness | -3762.64 | 7.58 | 0.01 |
|  | Season + Diet + Relatedness + Proximity | -3761.96 | 8.25 | 0.01 |
|  | - | -3713.73 | 56.48 | 0.00 |

To assess whether the effect of social connectedness on gut microbiome beta-diversity was driven by closely connected within-group dyads having very similar gut microbiomes, we repeated the analyses with between-group dyads only (N = 966 dyads). The full models and the social connectedness models were again the strongest supported models (Table 2). Beta-diversity was predicted by year and social connectedness, but not by diet and relatedness (Fig. 2).

### Socially structured OTUs

Our data set contained OTUs from 14 phyla, of which the most well-represented was Firmicutes, followed by Bacteroidetes, Spirochetes, and Verrucomicrobia (Supplementary Material Fig. 1). In each social group, at least 70% of the OTUs belonged to the phylum Firmicutes (Supplementary Material Fig. 1) and at least 50% of the OTUs belonged to the families Lachnospiraceae and Ruminococcaceae in the phylum Firmicutes (Supplementary Material Fig. 2).

Social connectedness predicted differences in abundances for 73 of the 396 OTUs in the 2009 data set (Moran’s I range: −0.27 – −0.14, all P < 0.05, Supplementary Material Table 1). The number of OTUs with a significant relationship to social connectedness was greater than expected in the phylum Firmicutes (N = 64, Hypergeometric test: P < 0. 001). The numbers of socially structured OTUs in the phyla Bacteroidetes (N=6), Planctomycetes (N = 1), Proteobacteria (N = 1) and Tenericutes (N = 1) were not greater than expected based on the total number of OTUs in these phyla (Hypergeometric tests, p > 0.050). The other phyla did not contain any socially structured OTUs. Four families had a higher than expected number of socially structured OTUs: Bacteroidaceae (N = 4), Lachnospiraceae (N = 20), Peptococcaceae 2 (N = 1), and Ruminococcaceae (N = 31). There was also a greater than expected number of socially structured OTUs in 14 of 34 genera (Fig. 4).

**Figure 4.**
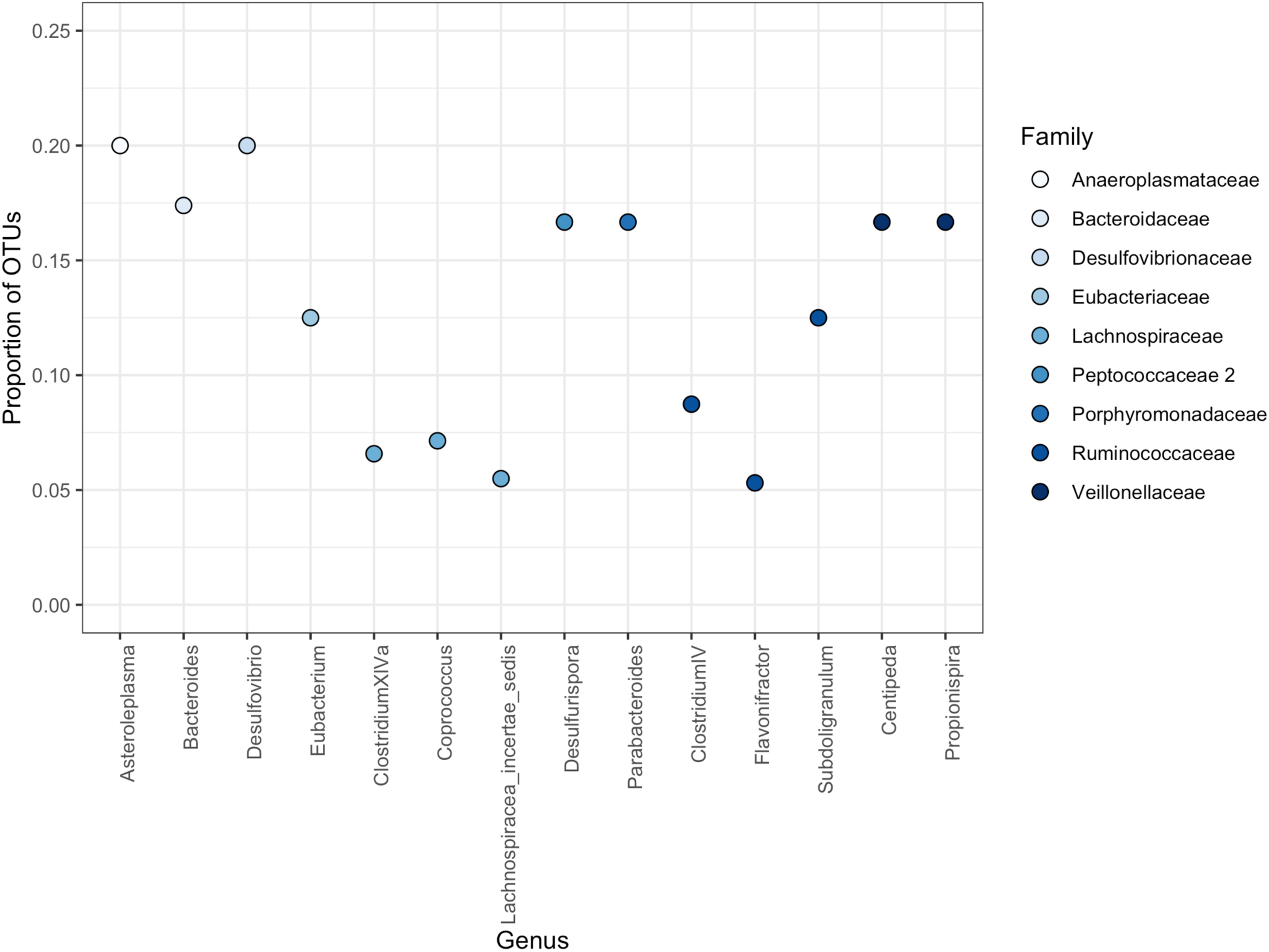
The proportion of Operational Taxonomic Units (OTUs) whose abundance was correlated with distance in the proximity network for genera that contained a higher than expected number of socially structured OTUs.

Social connectedness predicted differences in abundances for 1 of the 450 OTUs in the 2007 data set (Moran’s I range: −0.27 – −0.14, all p < 0.05), which belonged to the phylum Firmicutes, the family Lachnospiraceae, and the genus *Roseburia*. As a result, these taxa had a greater than expected number of socially structured OTUs (Hypergeometric tests, all p > 0.001).

## Discussion

The aim of this study was to investigate whether the increase in gut microbiome beta-diversity with home range separation in female colobus monkeys was best explained by diet, relatedness, or sociality. Distance in the proximity network was a better predictor than diet and relatedness, similar to findings in more social primates (Amato et al., 2017; Perofsky et al., 2017; Raulo et al., 2017; Tung et al., 2015). Although these previous studies suggest that strong social bonds within social groups drive between-group differences in the gut microbiome after ruling out the effects of relatedness and diet, this is the first report of a relationship between gut microbiome beta-diversity and social connectedness between individuals in different social groups. In contrast, gut microbiome dissimilarity between individuals residing in different social groups did not increase with grooming network distance in sifakas (Perofsky et al., 2017). These contrasting results may be due to the nature or frequency of the host population’s between-group interactions. Colobus monkeys sometimes engage in affiliative, sexual, and playful behaviours with non-group members (Supplemental Information; Sicotte & MacIntosh, 2004; Teichroeb, Marteinson, & Sicotte, 2005; Teichroeb et al., 2011), which differ from the almost exclusively aggressive nature of between-group encounters in some other taxa. Similar to these colobus monkeys, mountain gorillas (*Gorilla beringei beringei*) occasionally affiliate with members from other social groups (Forcina et al., 2019) and human foraging societies form extended social networks to optimize resource flow (Hamilton, Milne, Walker, Burger, & Brown, 2007). These extended networks could possibly affect their gut microbiome in similar ways as documented here in colobus monkeys.

To determine the consequences of such socially-mediated transmission, the first step is to determine which types of microbes are transmitted this way. The socially transmitted OTUs in this study included all taxa (family Porphyromonadaceae and genera *Parabacteroides* and *Coprococcus)* that diverged after a social group fission at our site (Goodfellow et al., 2019) and genera (*Bacteroides, Clostridium,* and *Roseburia*) that were transmitted via grooming and close proximity in howlers (Amato et al., 2017). The close match in socially transmitted taxa in howlers and colobus is not particularly surprising given both have a folivorous diet and low degree of terrestriality, which are factors that influence the gut microbiome (Perofsky et al., 2019). In contrast, the socially transmitted OTUs in our study did not overlap with those transmitted via grooming within social groups of baboons (Tung et al., 2015), despite the host species relatively close phylogenetic relationship. Recent findings show that host phylogeny has a stronger effect than diet on gut microbiome composition (Amato et al., 2019), and it is thus possible that while phylogeny has the strongest overall effect on the gut microbiome, the same gut microbial taxa are structured by sociality in primates with similar lifestyle.

We found that the majority of socially transmitted OTUs belonged to the most dominant families in our host population and other folivorous primates (Barelli et al., 2015; Perofsky et al., 2017), the families Lachnospiraceae and Ruminococcaceae in the phylum Firmicutes. These taxa are well-suited for breaking down hard-to-digest plant material (Biddle, Stewart, Blanchard, & Leschine, 2013), and it is therefore possible that socially transmitted gut microbes benefit hosts in terms of improved digestion of mature leaves, which make up the majority of the colobus diet (Saj & Sicotte, 2007). Several studies imply that socially-mediated transmission benefits the host. For example, Tung and colleagues (2015) suggest that the positive health and fitness effects that baboons accrue from forming close social bonds with group members are mediated by the gut microbiome. Our results support the notion that this may be the case in a wide range of gregarious species, including those with relatively low frequencies of social interactions. If social transmission sustains a healthy gut microbiome (as documented in Koch & Schmid-Hempel, 2011), it could provide an incentive for the formation and maintenance of social bonds within social groups (Lombardo, 2008). Our findings leave open the as-of-yet unexplored possibility that social transmission of microbes may even explain the occurrence of friendly between-group encounters, especially in the absence of limiting resources such as fertile females and important food sources.

The results of this paper ultimately lead us to an important outstanding question, which is how gut microbes are transmitted among animals that spend considerably less time grooming or in direct contact than other primates with socially mediated gut microbe transmission (Amato et al., 2017; Raulo et al., 2017; Tung et al., 2015). It might be that microbes are transmitted directly during the occasions we observed non-group members copulating, grooming, and playing. However, it could also be that the microbes are transmitted indirectly between hosts when they are touching shared surfaces within a certain time period (Münger et al., 2018). This reasoning is consistent with spatial proximity predicting the gut microbiome in other gregarious species with low frequencies of social behaviours like the Welsh Mountain ponies (*Equus ferus caballus*) (Antwis et al. 2018) and in more solitary species such as North American red squirrels (*Tamiasciurus hudsonicus*) (Ren et al., 2017) and gopher tortoise (*Gopherus polyphemus*) (Yuan et al., 2015). The occurrences of direct and indirect social transmission are difficult to tease apart when the two are correlated and when brief physical contact between extra-group members often go unnoticed, but carefully designed studies in the future may be able to address this question.

Finally, relatedness and dietary differences within a season were not good predictors of beta-diversity in comparison to social connectedness. In contrast, seasonal changes in diet may be associated with changes in the colobus gut microbiome, because beta-diversity was higher during the 2009 dry season when their diet was more diverse than during the 2007 rainy season when they ate mostly mature leaves. We will continue to investigate whether this seasonal dietary switch is linked to changes in the gut microbiome, as previously reported from other species inhabiting seasonal environments (Amato et al., 2015; Hicks et al., 2018; Orkin, Campos, et al., 2019; Smits et al., 2017; Springer et al., 2017). These authors concluded that gut microbiome dynamics determine nutrient uptake and is key for dietary flexibility (Amato et al., 2015; Hicks et al., 2018; Orkin, Campos, et al., 2019; Smits et al., 2017; Springer et al., 2017), while the potential three-way interaction between social, dietary, and gut microbial dynamics is still poorly understood. An interesting venue for further research is therefore to investigate whether the gut microbiomes of socially well-connected individuals map more quickly onto ecological changes, which could help them adjust to the rapidly changing environments that many wild animals inhabit today.

## Acknowledgements

We would like to thank two anonymous reviewers for their help with improving this manuscript; Keaton Stagaman for use of his custom pipeline; Fernando Campos, Teresa Holmes, and Robert Koranteng for field data collection; BFMS management committee, Ghana Wildlife Division, and the University of Calgary’s Life and Environmental Sciences Animal Care Committee for permission to conduct this study; Alberta Innovates Technology Futuresn (200*6*00343), American Society of Primatologists, Animal Behaviour Society, International Primatological Society, Leakey Foundation, Natural Sciences and Engineering Research Council of Canada, Sweden-America Foundation, Research Services at the University of Calgary, the National Institute of General Medical Sciences (P50GM098911) via the META Center for Systems Biology, and Wenner-Gren Foundation (8172) for funding.

## Data Accessibility Statement

All raw data are stored in the PaceLab database hosted by the University of Calgary. The 16S sequencing data will be uploaded to NCBI’s Short Read Archive. The data used for the analyses presented here will be uploaded to Dryad.

## Appendix

### Microbial taxa predicted to be structured by social connectedness

The last aim of this study was to investigate whether social connectedness was correlated with Operational Taxonomic Unit (OTU) abundances in certain taxa (Table A1) previously reported as structured by social relationships (Amato et al., 2017; Tung et al., 2015). We also expected social connectedness to be correlated with the abundances of three gut microbial taxa that diverged between two daughter groups after a group fission (DA and NP in Fig. A1), because we suspect that this pattern was driven by social network changes (Goodfellow et al., 2019).

**Figure A1.**
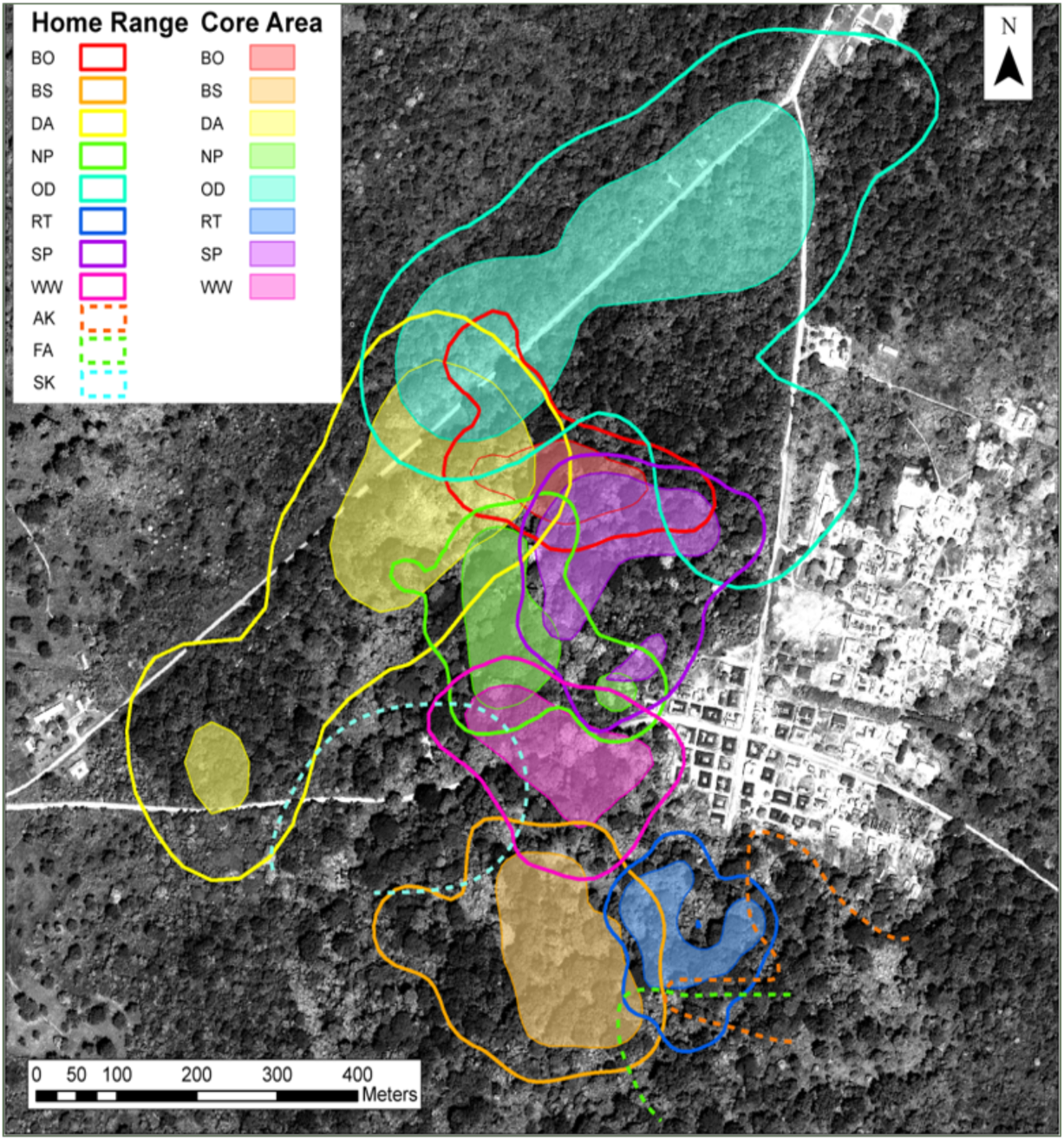
Home ranges and core areas for groups in the main forest fragment at Boabeng-Fiema, Ghana. Solid lines indicate home ranges of groups from which we collected behavioural data. Dashed lines indicate partial home ranges from other groups present in this forest.

**Table A1.**
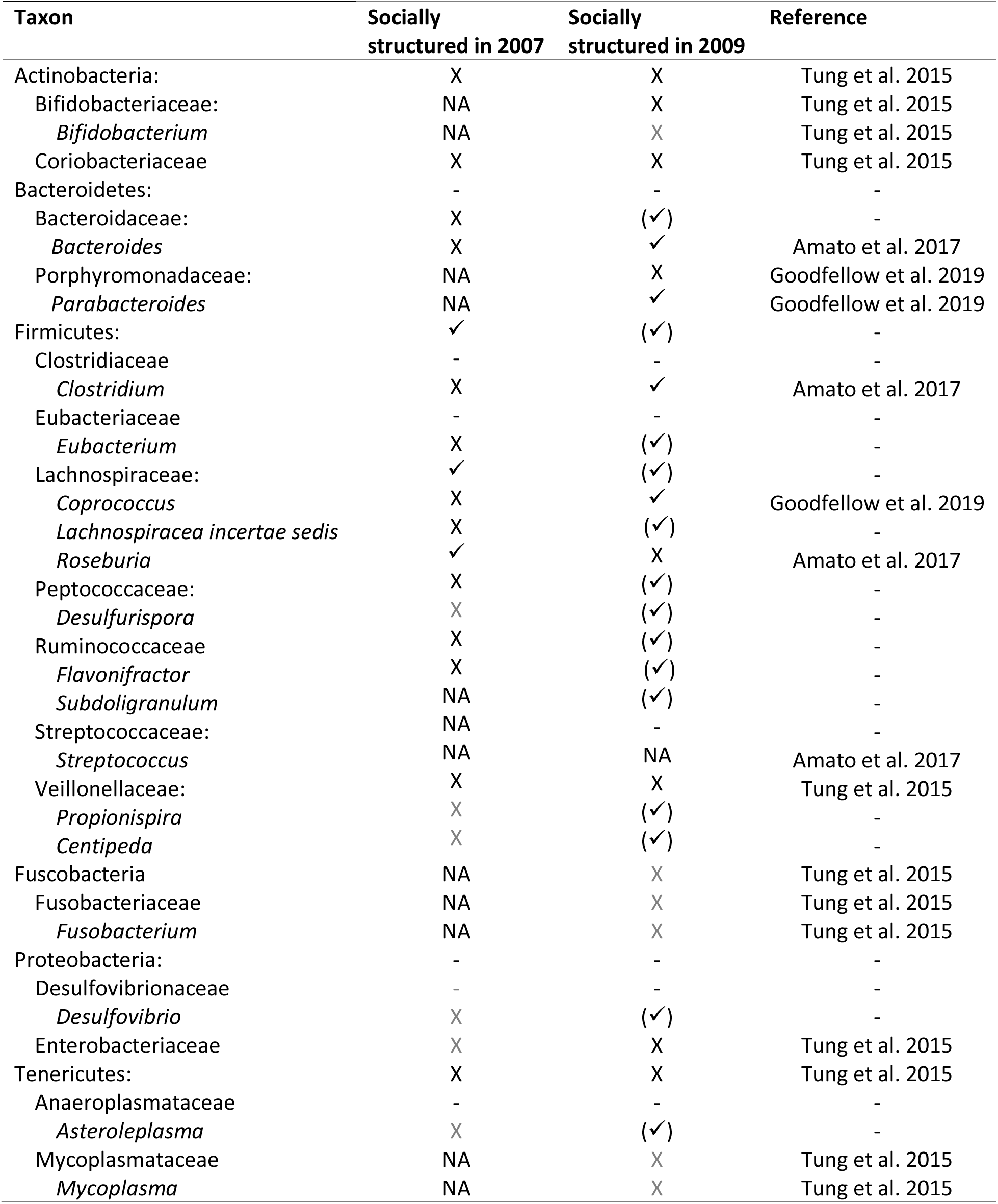
OTUs in these phyla, families, and genera are expected to be structured by sociality based on previous studies. Predictions were supported ✓; not supported X; no prediction made but structured in our data set (✓); or no prediction made and not structured in our data set (-). Grey text indicates rare taxa (N < 3 OTUs). NA denotes taxa not present in our data set.

| Taxon | Socially<br>structured in 2007 | Socially<br>structured in 2009 | Reference |
| --- | --- | --- | --- |
| Actinobacteria: | X | X | Tung et al. 2015 |
| Bifidobacteriaceae: | NA | X | Tung et al. 2015 |
| *Bifidobacterium* | NA | X | Tung et al. 2015 |
| Coriobacteriaceae | X | X | Tung et al. 2015 |
| Bacteroidetes: | - | - | - |
| Bacteroidaceae: | X | (✓) | - |
| *Bacteroides* | X | ✓ | Amato et al. 2017 |
| Porphyromonadaceae: | NA | X | Goodfellow et al. 2019 |
| *Parabacteroides* | NA | ✓ | Goodfellow et al. 2019 |
| Firmicutes: | ✓ | (✓) | - |
| Clostridiaceae | - | - | - |
| *Clostridium* | X | ✓ | Amato et al. 2017 |
| Eubacteriaceae | - | - | - |
| *Eubacterium* | X | (✓) | - |
| Lachnospiraceae: | ✓ | (✓) | - |
| *Coproccoccus* | X | ✓ | Goodfellow et al. 2019 |
| *Lachnospiraceae incertae sedis* | X | (✓) | - |
| *Roseburia* | ✓ | X | Amato et al. 2017 |
| Peptococcaceae: | X | (✓) | - |
| *Desulfurispora* | X | (✓) | - |
| Ruminococcaceae | X | (✓) | - |
| *Flavonifractor* | X | (✓) | - |
| *Subdoligranulum* | NA | (✓) | - |
| Streptococcaceae: | NA | - | - |
| *Streptococcus* | NA | NA | Amato et al. 2017 |
| Veillonellaceae: | X | X | Tung et al. 2015 |
| *Propionispira* | X | (✓) | - |
| *Centipeda* | X | (✓) | - |
| Fusobacteria | NA | X | Tung et al. 2015 |
| Fusobacteriaceae | NA | X | Tung et al. 2015 |
| *Fusobacterium* | NA | X | Tung et al. 2015 |
| Proteobacteria: | - | - | - |
| Desulfovibrionaceae | - | - | - |
| *Desulfovibrio* | X | (✓) | - |
| Enterobacteriaceae | X | X | Tung et al. 2015 |
| Tenericutes: | X | X | Tung et al. 2015 |
| Anaeroplasmataceae | - | - | - |
| *Asteroleplasma* | X | (✓) | - |
| Mycoplasmataceae | NA | X | Tung et al. 2015 |
| *Mycoplasma* | NA | X | Tung et al. 2015 |

### DNA extraction, amplification, and sequencing protocols for the gut microbiome analysis

We extracted DNA from 200 μl of sample using QIAmp DNA stool extraction protocol the following modifications. Step 2: Added 50 μl Proteinase K with overnight lysis before proceeding to step 3. Step 4: Pipetted all of the supernatant. Step 5: Used half of the InhibitEX tablet. Step 6: Centrifuged for 5 minutes. Step 9: Added 4ul RNAse and vortexed for 15 seconds. Step 19: Used 50 μl Buffer AE and incubated at 10 minutes. Step 20: Pipetted the same 50 μl of buffer AE back onto filter and incubated at room temperature for 15 minutes. Centrifuged at full speed for 2 minutes. Our DNA extraction protocol did not include a bead-beating step, which could bias against lysis-resistant taxa such as Gram-positive and spore-forming bacteria that are less likely to be dependent on direct social contact for transmission between hosts because they can survive for prolonged periods outside the host (Pollock, Glendinning, Wisedchanwet, & Watson, 2018; Yuan, Cohen, Ravel, Abdo, & Forney, 2012).

We determined the concentration of the extracts using Qubit dsDNA BR Assay Kit (Invitrogen) and diluted products to 2nM for downstream reactions. We amplified the bacterial v4 region of the 16S ribosomal RNA gene using the following 515F and 806R primers containing 5’ Illumina adapter tails and dual indexing barcodes:

515F 5’ AATGATACGGCGACCACCGAGATCTACACTAGATCGCTATGGTAATTGTGTGCCAGCMGCCGCGGTAA

806R 5’ CAAGCAGAAGACGGCATACGAGATTCACCTAGAGTCAGTCAGCCGGACTACHVGGGTWTCTAAT.

We set the PCRs with 12.5 μl NEB Q5 Hot start 2x Master mix, 1.25 μl 10uM Primer mix, 1 μl template DNA, and 10.25 μl MoBio certified DNA free water and used the following cycling protocol: 98 degrees for 30 seconds (1x) followed by 98 degrees for 10 seconds, 61 degrees for 20 seconds, and 72 degrees for 20 seconds (20x), followed by 72 degrees for 2 minutes and 4 degrees. The amplification products were cleaned up using Ampure XP beads and normalized into a final pool with an Eppendorf liquid handling robot. Libraries were sequenced as part of a 150bp paired-end sequencing run on the Illumina NextSeq platform following the manufacturer’s protocol.

We used a custom pipeline that contained the following steps: joining pair-end reads; removing low-quality and chimeric reads; dereplication and dropping unique reads with low abundance; clustering OTUs; making OTU table; alignment; building a reference tree; and taxon assignment using FLASH (Magoc & Salzberg, 2011), the FASTX Toolkit (Hannon Lab, 2010), and the USEARCH pipeline (Edgar, 2010). See https://github.com/kstagaman/Process_16S and Goodfellow et al. (2019) for further details. We performed *de novo* OTU picking in UCLUST (Edgar, 2010), and sequences with 97% overlap were defined as belonging to the same bacterial Operational Taxonomic Unit (OTU). To guard against sequencing errors, we filtered out OTU’s with a frequency lower than 0.00005 as recommended (Bokulich et al., 2012).

### Variation in predictor and outcome variables

Of the females included in the analyses with behavioural predictor variables (Table A2), dietary dissimilarity (i.e., Sørensen diversity index) varied from 0 to 1, dissimilarity in relatedness calculated as their *R*-value subtracted from 1 ranged from 0.31 to 1, and social connectedness (i.e., inversed path length or Geodesic distance in the 1-meter proximity network) varied from 0 to 1 where 0 represents unconnected dyads (Fig. A2).

In our full data set, mean unweighted Unifrac distances within the same season and year was 0.052 ± 0.004 for samples collected from the same individual (N = 4 samples) and 0.205 ± 0.042 for samples collected from different individuals within the same season and year (N = 61 samples). The low amount of within-individual variation in comparison to the between-individual variation suggests that one sample per individual is representative of its gut microbiome during that season and sufficient for analysis of beta-diversity. Furthermore, the beta-diversity of matched samples from the same adult female in the wet season 2007 and the dry season 2009 (N = 22 samples from 11 females) was lower (0.152 ± 0.031) than the female’s mean beta-diversity with samples from a different female and year (0.183 ± 0.021) for all but one female, and there was a significant difference in beta-diversity between samples from the same versus different females in this sample (N = 11 females, Wilcoxon signed rank test, p < 0.001).

**Table A2.**
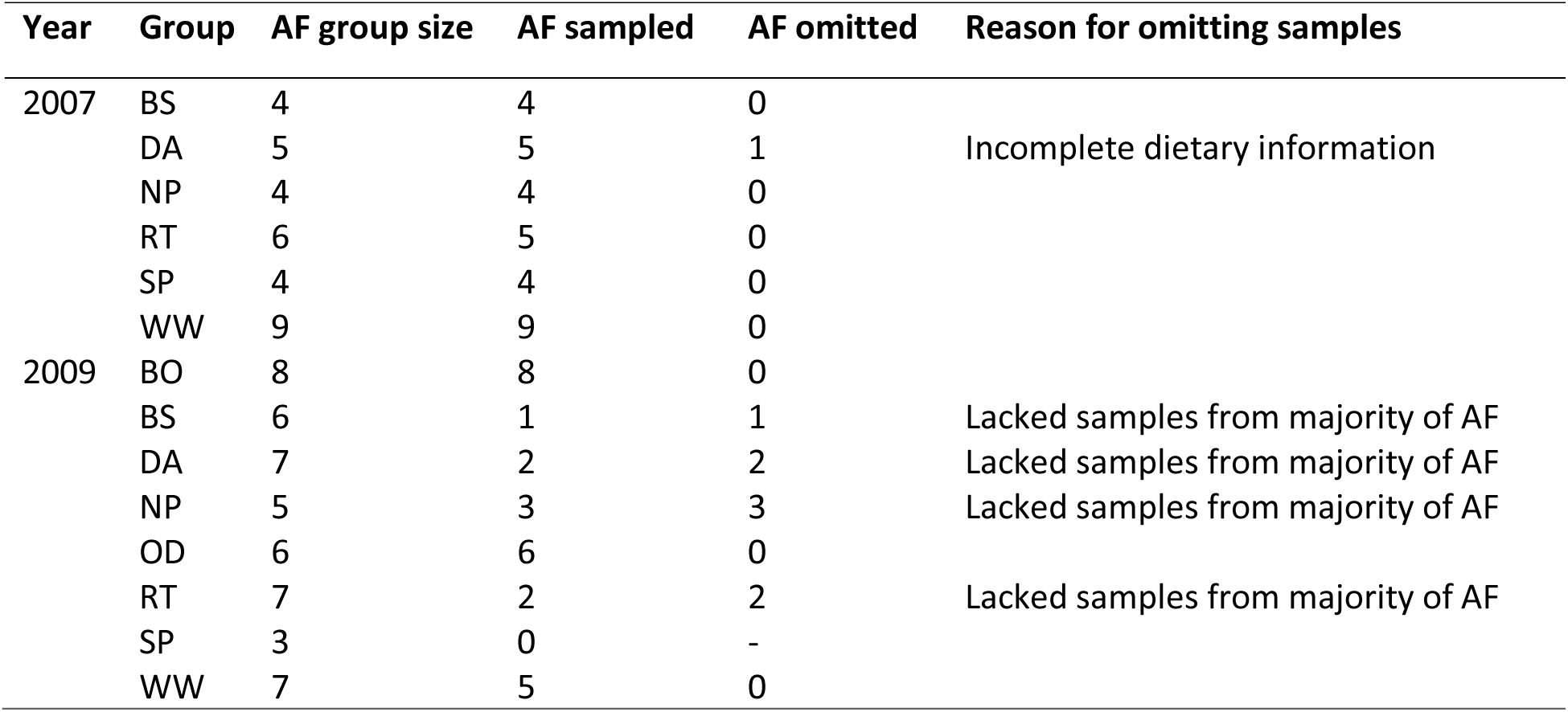
Number of adult females (AF) present, sampled, and omitted from data analyses with behavioural predictor variables.

**Figure A2.**
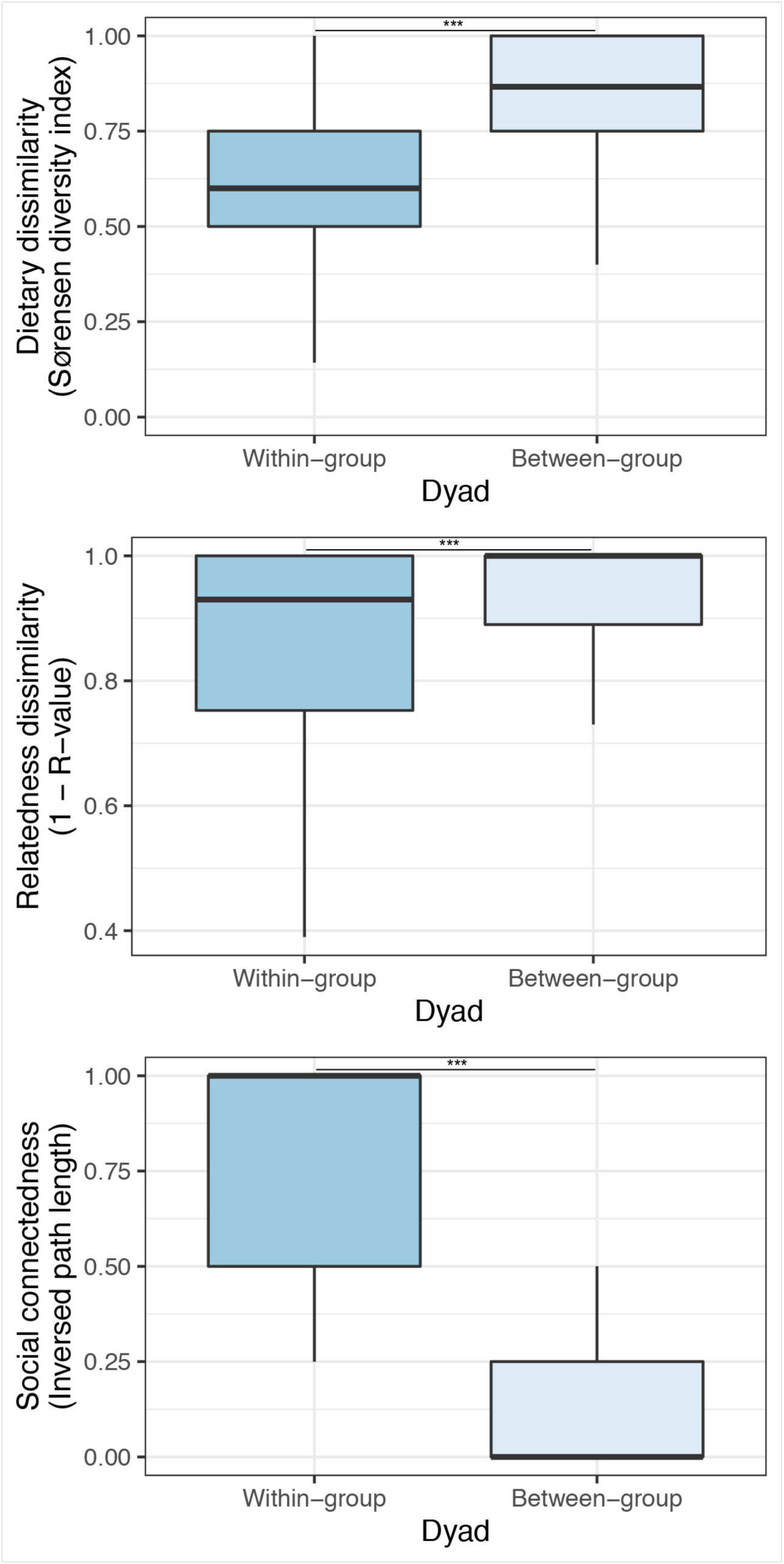
Dietary dissimilarity, relatedness dissimilarity, and social connectedness were lower for within-group than between-group female-female dyads (Wilcoxon signed rank tests, N = 88 samples, all p < 0.001).

